# Unsupervised spike sorting for large scale, high density multielectrode arrays

**DOI:** 10.1101/048645

**Authors:** Gerrit Hilgen, Martino Sorbaro, Sahar Pirmoradian, Jens-Oliver Muthmann, Ibolya E. Kepiro, Simona Ullo, Cesar Juarez Ramirez, Albert Puente Encinas, Alessandro Maccione, Luca Berdondini, Vittorio Murino, Diego Sona, Francesca Cella Zanacchi, Evelyne Sernagor, Matthias H. Hennig

## Abstract

A new method for automated spike sorting for recordings with high density, large scale multielectrode arrays is presented. Exploiting the dense sampling of single neurons by multiple electrodes, we obtain an efficient, low-dimensional representation of detected spikes consisting of estimated spatial spike locations and dominant spike shape features, which enables fast and reliable clustering into single units. Millions of events can be sorted in minutes, and the method is parallelized and scales better than quadratically with the number of detected spikes. We demonstrate this method using recordings with a 4,096 channel array, and present validation based on anatomical imaging, optogenetic stimulation and model-based quality control. A comparison with semi-automated, shape-based spike sorting exposes significant limitations of conventional methods. Our analysis shows that it is feasible to reliably isolate the activity of hundreds to thousands of neurons in a single recording, and that dense, multi-channel probes substantially aid reliable spike sorting.

## Introduction

Large scale, dense probes and arrays and planar multielectrode arrays enable extracellular recordings of thousands of neurons simultaneously (Eversmann et al., 2003; Berdondini et al., 2005; Hutzler et al., 2006; Frey et al., 2010; Ballini et al., 2014; Maccione et al., 2014; Müller et al., 2015; Obien et al., 2015). Exploiting such data requires the reliable isolation of extracellularly recorded spikes generated by single neurons, spike sorting, a computationally costly task that is difficult to scale up to large numbers of recording channels (Rey et al., 2015). For conventional devices with up to tens of recording channels, a typical workflow consists of initial event detection, followed by semi-automated clustering based on spike waveform differences, and manual inspection and refinement of the proposed event assignments. If the recording channels are sufficiently well separated, there is no or little overlap between their signals, and spike sorting can be performed by clustering a low-dimensional representation of spike shapes (e.g. Lewicki, 1998; Harris et al., 2000; Quiroga et al., 2004).

This approach, however, is inappropriate for dense, large scale recordings for two rather obvious reasons. First, on dense arrays spike sorting becomes a very complex assignment problem, since not only multiple neurons contribute to the compound signal recorded on individual channels, but each neuron’s spikes is also recorded by several neighboring channels simultaneously (Prentice et al., 2011; Rossant et al., 2016). Events are thus described by multiple waveforms and their locations, with an exponential number of potential assignments that can only be tackled using approximate algorithms. Second, the sheer size of the data sets makes extensive manual intervention impractical, hence as much of the process as possible, including quality control, should be automated.

In addition, the measured signals in dense recordings differ from conventional recordings in a rather subtle way. Much of the variability in spike shapes is due to measuring them at different positions relative to the neuron. In conventional recordings, relatively small signals are measured using large electrodes averaging currents originating from different parts of the neuron. High density MEAs with small electrode detect primarily strong currents at the axon initial segment (AIS) of the neurons. In other words, the mechanism for generating action potentials is represented with a higher weight in the measured signals, leading, in terms of signal power, to a lower variability in the measured spike shapes. Existing solutions addressing these issues, which have been demonstrated on data from hundreds of channels, are template matching methods (Prentice et al., 2011; Marre et al., 2012), and the elimination of the effect of uninformative spike features such that fitting of a mixture model becomes feasible for large data sets (Rossant et al., 2016).

Here we present a new, very fast and fully automated method for spike sorting. Dense sampling enabled us to obtain a rough estimate of a source location for each detected event (Muthmann et al., 2015). Spikes detected this way tend to form dense, spatially separated clusters, which, as we show using optogenetic stimulation and confocal imaging, originate from single neurons. In addition, for each event an average waveform is obtained, with noise reduced by signal interpolation. Shape features extracted from this waveform are then combined with spatial locations, such that the clustering problem is reduced to finding local density peaks in few dimensions.

We demonstrate this method using light responses in the mouse retina and spontaneous activity in cell cultures recorded with a 4,096 channel array. To evaluate the quality of the sorting, post hoc analysis is performed on the clustered data, which allows largely unsupervised rejection of poorly clustered units, and highlights potentially problematic cases for further inspection. A direct comparison with conventional spike sorting also exposes severe, and hard to detect, limitations of the latter. A parallelized implementation of this method, which is capable of sorting millions of spikes within a few minutes on a fast workstation, as well as a tool for data visualization, can be downloaded at https://github.com/martinosorb/herding-spikes.

## Results

### Spatial spike localization

Figure 1A illustrates a typical recording setup, with a flattened retina placed on the array. In a first step, spikes were detected using threshold-based method that exploits dense sampling to improve detection performance, and assigns each spike an estimated location based on the barycenter of the spatial signal profile (Muthmann et al., 2015). This procedure yields spatio-temporal event maps, where each event is identified by a time stamp, two spatial coordinates and a single, interpolated waveform. The resulting spatial activity maps provided a higher spatial resolution for spike locations than given by the electrode arrangement of the MEA (Figure 1B). As expected for signals originating from distinct sources, spikes were found in dense clusters surrounded by areas of low event density. The relationship between recorded signals and spike locations is illustrated in Figure 1C (magnified part of Figure 1B). The estimated locations of several spikes are indicated by circles, and the corresponding raw data segments highlighted in the traces from nearby electrodes. These examples show how spike locations are related to the spatial decay of the voltage peaks, and that the decay was sufficiently wide to estimate their peak locations on the array (Figure 1D, cf. Pettersen & Einevoll 2008; Linden et al. 2011; Mechler et al. 2011).

**Figure 1:**
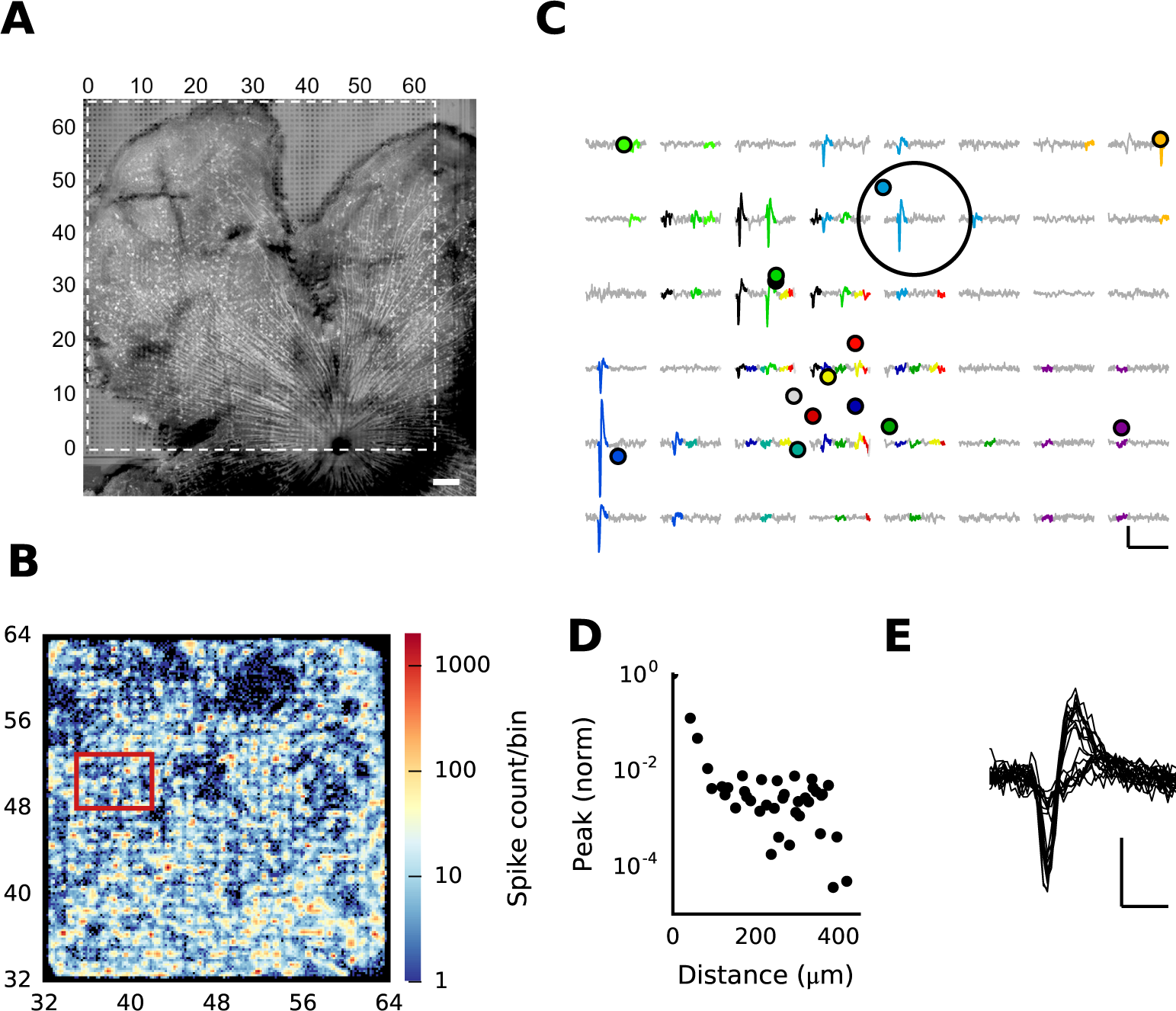
Spatial event localization reveals isolated spike clusters. (A) Confocal image of a Thy1-ChR2-YFP retina expressing yellow fluorescent protein under the Thy1 promoter, placed on the array for recording. Electrodes can be seen as small squares in areas not covered by the retina. The active area of the array is indicated by dashed lines. Scale bar is 200 *μ*m. (B) The activity map of a quarter of the array after spatial event localization. Spike counts shown as density plot, spatially binned with 8.4 *μ*m resolution. This reveals distinct clusters in space, presumably originating from individual neurons. The data in panels B-E are from the same recording acquired at 24 kHz on 32x32 channels, panel A shows a different preparation. (C) Examples of several detected events (rectangle in B), shown at their estimated locations (colored circles), and the corresponding episodes in the raw data (colored traces). Scale bars are 5 ms and 200 *μ*V. (D) Average peak signal decay for detected events as a function of distance. On average, a significant signal is detectable in an area of 100 *μ*m around the spike peak location. Note that this plot is based on signal peaks at the spike time ±2 recording frames, so signals beyond 200 *μ*m reflect primarily noise. (E) Twenty randomly selected spike shapes for events localized within the area marked by the large circle in panel C, indicating the presence of signals from at least two different neurons at this location. Scale bars are 5 ms and 200 *μ*V.

Thus, on dense MEAs event locations provide a compact summary of the spatial activity footprint for each spike. An inspection of waveforms, however, reveals the presence of multiple units in small areas (Figure 1E). This shows that clustering spatial locations alone is insufficient for reliable single unit isolation, as also shown in earlier work (Prentice et al., 2011).

### Combined spatial and shape-basedl clustering

Next, spikes are clustered using a combination of their estimated locations and their dominant waveform features, extracted via PCA. Together, these two feature sets provide a complementary, compact description of the events. The location estimate is an effective way of summarizing the spatial footprint each spike leaves on the array, while waveforms enable the separation of spatially overlapping sources, and they remove ambiguities at spatial cluster boundaries.

The Mean Shift algorithm was used for clustering these data, with the number of clusters automatically determined and is controlled by a single scale parameter (Comaniciu & Meer, 2002). Clusters are formed by moving spikes along density gradients, and augmented by local differences in spike waveforms. We found that including the first two principal components was sufficient to successfully isolate single units. This reduces the high dimensional assignment problem to four dimensional clustering, which can be performed in minutes for millions of events on a fast computer. In addition to the scale parameter, this method only requires a mixing coefficient for the shape information as a second parameter.

Figure 2A-C shows the result of clustering a recording from 1,024 channels at 24 kHz sampling rate, which had about 440,000 spikes that were separated into 1,600 units. Cluster sizes ranged from tens of spikes to several thousands, corresponding to firing rates ranging from 0.1 to 30 Hz (circle areas in Figure 2A indicate the firing rate of each unit). A magnified view of a subset of clusters, together with the average spike waveforms, shows that units may indeed spatially overlap, but are separated by their waveform features (Figure 2B). Overall, units with clearly bi-phasic and large amplitude waveforms tend to form the more spatially coherent clusters, while smaller events are spatially more spread out (compare average waveforms for the clusters).

**Figure 2:**
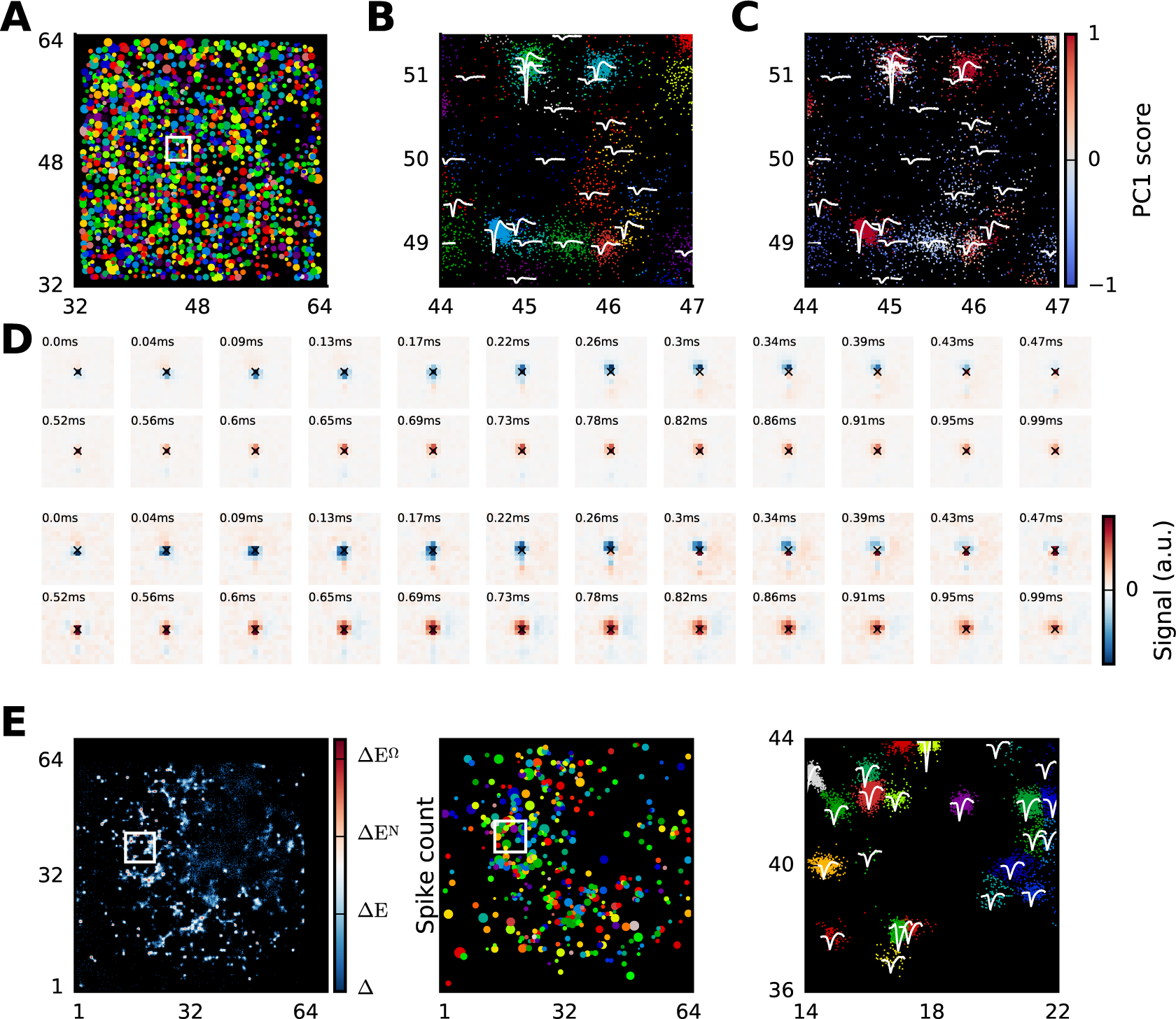
Illustration of clustered spike data. (A) Overview of all single units obtained by clustering a recording from the retina (same data as in Figure 1, acquired at 24 kHz), shown as circles at their estimated locations in array coordinates. Circle areas are proportional to firing rates. (B) Magnified view of a group of units (area in white rectangle in panel A), showing a subset of spikes at their estimated locations (dots, colored by unit membership, same colors as A), and the average waveform associated with each unit. (C) As B, but with spike colors encoding the magnitude of the spike waveform projection along the first principal component (PCl score). Higher scores represent bi-phasic waveforms, and low scores weak deflections without a clear bi-phasic shape. (D) Electrical images for two units, negative signals relative to baseline are colored in blue, and positive signals in red. The cross indicates the centroid of the spike locations. Each square represents one electrode, l5 x l5 (0.63 x 0.63 mm) electrodes are shown. Axonal propagation can be seen, moving downwards towards the optic disk. (E) Clustered recording from a hippocampal culture. The panels show raw spike counts (left), all units obtained during the clustering step (middle), and a magnified view of a small area of the MEA showing individual spikes and average unit waveforms (right). This recording was acquired with 4,096 channels at 7 kHz sampling rate, and a waveform classifier was used to remove noise prior to clustering (see Supplementary Material, Figure Sl).

The units with small waveforms originate from neurons that left weak signals on the MEA, but that were nevertheless detected as we generally used a low threshold during the detection step to avoid false negatives. Such units may contain an incomplete spike records, or contain signals from multiple neurons that could not be well separated. The first PC projection (PC1) for the events is a good indicator of their biphasic character, and using the convention that positive values always coincide with more biphasic waveforms, this measure can be used to (de-)select units for subsequent analysis (Figure 2C). A similar method, which we used for all recordings performed at lower sampling rates (7 kHz, 4,096 channels), was to train and employ a classifier to pre-select valid spikes prior to clustering based on salient waveform features (see Supplementary Material, Figure S1). This method can remove noise more reliably than simply increasing the higher detection threshold or using other pre-specified criteria on spike shapes, as the classifier is better adjusted to the specific recording conditions. Importantly however, this step is not required for sampling rates of >10 kHz.

As a first assessment of the clustered units, we generated electrical images for individual units (two examples are shown in Figure 2D). These images provide a spatio-temporal representation of the raw signal recorded by the MEA around the time of spiking, and is generated as a spike-triggered average of the signal on each electrode. Out of 406 units with at least 100 spikes inspected in this way, all but one had an estimated location within 40 *μ*m of the electrode that contained the peak signal (less than one electrode diameter; median distance 9.7 *μ*m), indicating that units are indeed well aligned with their spatio-temporal electrical footprint. Furthermore, the two examples shown in Figure 2D reveal that the recordings were of sufficient detail to isolate axonal propagation. In both cases, a separate, weak positive peak, followed by a negative peak, can be seen traveling downwards (towards the optic disk). Since these events peak within less than 100 *μ*s of the main signal, they are not detected as separate events, but instead introduce a small bias on the location estimates during spike localization.

We also tested our method on activity recorded from cultured hippocampal neurons. The example in Figure 2E illustrates that isolation of single units is also feasible for these preparations, although here the spike localization was less precise than in the retina. We attribute this to the effect of a larger effective conductivity in the space above the electrodes, which is expected to reduce recorded signal amplitudes, which in turn increases the influence of noise on localization (Ness et al., 2015). In contrast, this conductivity is likely much lower for the approximately 200 μm thick retina, leading to larger and more precisely localizable signals. Ness et al. (2015) show that in this case even small MEA-tissue gaps strongly reduce the signal recorded amplitude, a likely explanation for the clear, sharp boundaries between areas with and without recorded spikes. Yet as also observed for the retina, spikes in cultures were typically spatially well clustered, and waveform differences had sufficient detail to allow separation of overlapping units (see Figure 2E, right panel, for examples).

### Waveform features are essential for reliable clustering

To assess the importance of including waveform features in the clustering process, and the role of the mixing coefficient *α* that determines the weight of waveform features relative to locations during clustering, we compared the correlations between all waveforms within each unit with cross-correlations of waveforms between this unit and its closest neighbor, or all nearby spikes within a radius of 42*μm* (electrode pitch). A well sorted unit is expected to have high within-correlations, and smaller cross-correlations. Three examples of this analysis are shown in Figure 3A-C. The first illustrates a case where spatial clustering alone was sufficient to isolate a unit (Figure 3A). The correlations after clustering spatial locations alone (*α* = 0) are very similar to those obtained when waveforms are added (*α* = 0.3), although in the latter case a few spikes were re-assigned based on their waveform features. In contrast, in the other two examples two clearly distinct units were spatially overlapping, and could therefore only be separated when waveform features were included (Figure 3B,C). In these cases, increasing *α* causes a substantial increase in self-correlation. When nearby events have sufficiently distinct waveforms, this also leads to a reduction in cross-correlations with events in other nearby units (Figure 3B), although this behavior is variable and differs if, for instance, spatially well isolated nearby units have similar waveforms (Figure 3C).

**Figure 3:**
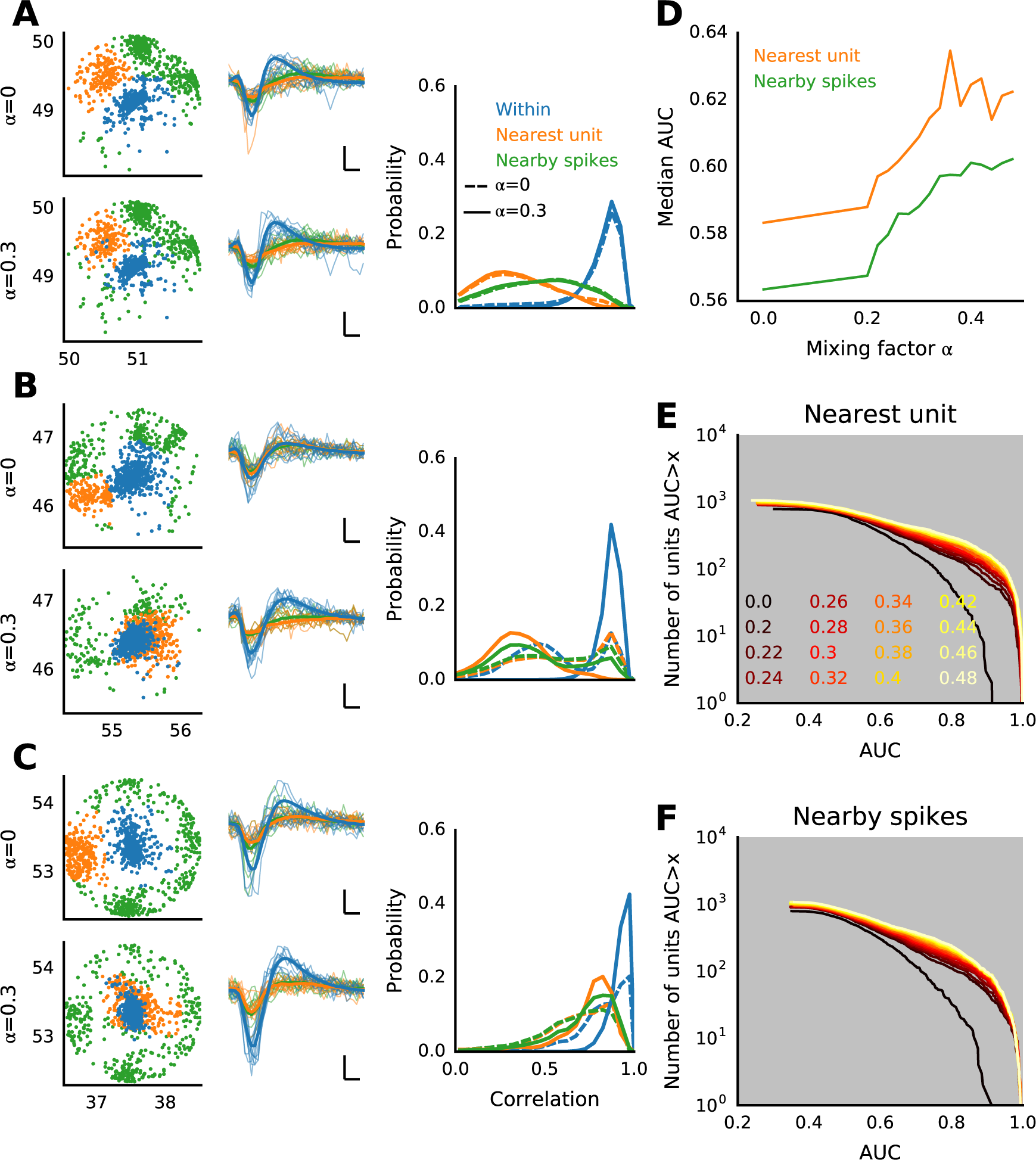
Waveform correlations demonstrate improved clustering for combined event locations and waveform features. All data is from the same experiment as in Figure 1, acquired at 24 kHz. (A-C) Three examples comparing the same unit obtained using spatial clustering alone (with mixing coefficient *α* = 0) with combined clustering based on combined event locations and waveform features (*α* = 0.3). Each panel shows event locations (left), example (thin lines) and average (thick lines) waveforms (middle, scale bars indicate 0.2 ms and l00 *μV*), and normalized distributions of waveform correlations (right, dashed lines: *α* = 0, solid lines: *α* = 0.3). The selected unit is colored in blue (within), the nearest unit in orange, and the remaining events within a radius of 42 *μm* of the target unit location in green (nearby spikes, these also include the spikes of the nearest unit). (D) Median area under ROC curves (AUC) for all units, quantifying the overlap between the normalized distributions of waveform correlations for each unit and with spatially neighboring events, as a function of the mixing coefficient *α*. The comparison was either done with the spatially closest unit (orange), or with all neighboring spikes (green). (D-E) Full distributions of AUC values for all units for different values of *α*.

To quantify the separability of these distributions, we computed, for each unit, the area under the receiver operating characteristic (ROC) curves (AUC), constructed from the distributions of self-correlations and correlations with events in the nearest unit (Figure 3E), or all neighboring events (Figure 3F). The area was calculated as the integral of the area spanned by the probability of findings a self-correlation above a sliding threshold, as a function of the probability of finding a cross-correlation above this threshold (true positives versus false positives), such that a value of 1 corresponds to perfectly separated distributions, while zero indicates full overlap.

The median AUC for all units in one recording increases as *α* is increased, before plateauing at values about *α*≈0.4 (Figure 3D). This indicates the combined features yield an overall improved separation of single units. The AUC distributions show that this effect is indeed substantial (Figure 3E,F). While spatial clustering alone only yielded three (out of 788 units with >100 spikes) units with AUC>0.9 when compared to events from its closest neighbor, this increases to 130 (out of 956 units with >100 spikes) for *α* = 0.32. This number rapidly increases when *α*≈0.25, and plateaus for larger values, indicating that the precise choice of this parameter is not critical. It is important to note that while high AUC values indicate a well isolated unit based on waveform features alone, hence units with small AUC should not be rejected as they may still be spatially well isolated.

In summary, waveform features help both to refine existing units found by spatial clustering, and to separate spatially overlapping units. The analysis further shows that event locations and waveforms indeed provide complementary information, and provide an effective way of summarizing the key features of the spatio-temporal footprint left by spikes on the array.

### Validation with optogenetics and anatomical imaging

To test whether the detected units indeed correspond to single neurons, we used Thy1-ChR2-YFP retinas (see Methods) expressing Channelrhodopsin-2 (ChR2), a light-gated cation channel, under the Thy1 promoter in about half of all RGCs (Raymond et al., 2008). This allowed us to stimulate spiking exclusively in a subset of visually identifiable RGCs, to clearly establish correlates between single spike-sorted units and individual RGCs.

We first compared the photoreceptor-driven activity recorded during normal light stimulation (irradiance 4 *μ*W/cm^2^, full field flashes at 0.5 Hz) with recordings obtained when light responses originating from the photoreceptor-bipolar-ganglion cell pathway are blocked with 20 *μ*M DNQX and L-AP4, and at a maximum irradiance of 0.87 mW/cm^2^ to selectively evoke ChR2 mediated spikes (Figure 4A). The activity maps show that only a subset of all RGCs responded to optogenetic stimulation (Figure 4A, compare top and middle plots). For these data, we found 375 units with a firing rate of at least 0.5 Hz during photoreceptor-driven light stimulation, but only 254 units during direct stimulation of ChR2-expressing RGCs. In addition, 77 units were significantly less active during light stimulation than during ChR2 stimulation, presumably neurons unresponsive to our light stimulus but nevertheless expressing ChR2. To assess the responsiveness of each unit to ChR2 activation, which was triggered by a random sequence of light flashes, we determined the correlation of an individual unit’s activity with the overall population activity (Figure 4A, bottom). Most, but not all highly active units during ChR2 stimulation also showed higher correlation, hence in some cases intrinsic spontaneous activity, which could not be blocked, was also present. About 40% of all detected units had a correlation larger than 40%, indicating that they were well activated.

**Figure 4:**
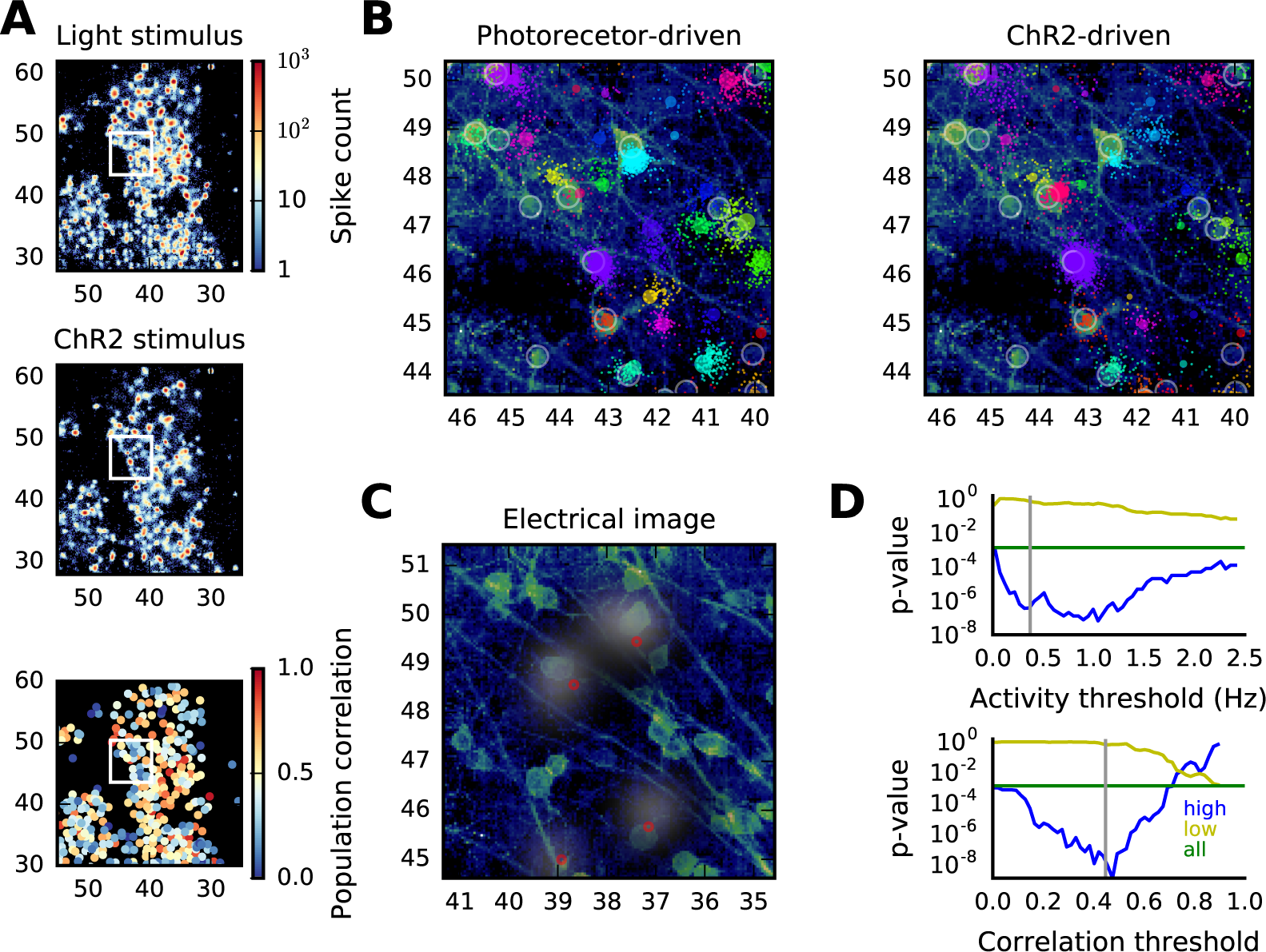
Comparison of optogenetically evoked spikes with anatomical imaging. (A) Activity maps obtained during light photoreceptor stimulation (top) and ChR2-expressing RGC stimulation under blockade of the glutamatergic pathway from photoreceptors to RGCs (middle). The bottom graph shows the correlation of the activity of each unit with the overall ChR2-driven population activity, which quantifies the responsiveness to optogenetic stimulation. (B) Alignment of neural activity with a confocal image. Individual spikes are shown as dots, colored according to unit membership (note only a subset of all recorded spikes are shown for clarity). Annotated somata are highlighted by circles, and unit’s centroids as colored circles, with areas proportional to the spike rate. (C) A different imaged area, with superimposed electrical images of four selected units. Cluster centroids are indicated a red circles. (D) The distribution of spatial distances between each unit and its closest soma is significantly different from randomness. The one-tailed Kolmogorov-Smirnov test shows incompatibility with the distribution obtained by assuming somata and units are unrelated (*p* = 0.00l, green line). When the units are separated into two sets according to activity level (top) and population correlation (bottom), the effect is strongest for highly active/highly correlated units (blue), while weakly active/correlated units are randomly distributed (yellow). The gray line indicates the threshold value for which the two sets have the same number of units. Data in these graphs summarize an imaged area of 0.78 mm^2^.

Next we co-localized the activity with confocal micrographs showing the YFP labeled neurons (Figure 4B). In total, we analyzed an area of 0.78 mm^2^, where 195 somata were manually annotated, and 211 units were detected (note that the spike sorting was performed on all recordings together). An example of the alignment of activity and anatomical image is shown in Figure 4B, for activity obtained during light stimulation (left) and ChR2 activation (right). While some units were clearly active for both stimuli, others had more spikes in only one condition. Importantly, all units with significant activity during ChR2 stimulation were closely co-localized with a soma. Similarly, a tight co-localization between the neurons and electrical images generated from the raw traces was observed (Figure 4C).

We now asked whether labelled somas and localised units where close to each other in a statistically significant way. For every unit, we computed the distance to the closest soma. If units and somata were randomly distributed, the probability of a distance *r* would be 2*πnre*^−*πnr*^2^^, where *n* is the density of somata (Chandrasekhar, 1943). We compared the distribution of 198 distances to this null model using a one-tailed Kolmogorov-Smirnov test (Figure 4D), which confirmed that the distances are significantly smaller than predicted by the random model. Next, to account for spontaneously active neurons, we applied the test after separating the units in two groups according to their activity level or population correlation, varying the threshold that separates the two sets. This showed that the locations of the less active and less correlated units locations are compatible with a random distribution, while the more active and more highly correlated units are significantly closer to their anatomical counterparts.

### Model based validation and quality control

The list of units provided by the clustering step contained assignments for all putative spike events detected in the raw data, regardless of their signal quality. As pointed out above, detection was performed with a low threshold to minimize false negatives. Hence a certain fraction of the units is expected to contain ambiguities the clustering algorithm cannot fully resolve. For instance, the localization error is typically larger for spikes with small amplitudes (Muthmann et al., 2015), hence it may not be possible to spatially cluster these events reliably.

To assess the quality of the cluster assignments, and to automatically reject poorly separated units, we followed an approach proposed by Hill et al. (2011). Under the assumption that the features of a unit (the spike locations and waveform features) can be described by a multivariate normal distribution, a comparison of the clusters assignments with those predicted by a Gaussian mixture model provides an estimate of the classification performance. This was done by investigating each unit in turn, including all nearby units which could interfere with the unit in question, and fitting a six-dimensional mixture model with the number of Gaussian given by the number of units (see Methods). We included four PCA dimensions to ensure the model best exploits all available waveform features, while ensuring convergence, which was impaired when more dimensions were included that typically had little or no structure that could be exploited to improve the fit. To evaluate the relevance of spatial locations and waveform features for clustering, the model was also fit to each of these features separately.

The model comparison produces a confusion matrix with the estimated number of false positives and negatives for each unit, which is then summarized into a single measure (F-score). Two typical outcomes of this procedure are illustrated in Figure 5A and B, for relatively crowded areas on the array. The first example was a unit with a distinct waveform (blue), which had four neighbors within one electrode radius (Figure 5A). In this case, the blue unit was already well isolated based on waveform features alone (PCA, F-score = 0.97), but not when only spike locations were considered (X/Y, F-score = 0.68). The combination of locations and waveforms did not yield further improvement, but somewhat helped to isolate its neighbors, primarily based on their spike locations. Note that separate F-scores were also computed for the neighboring units, using different sets of nearby units. The second example shows five spatially well separated units, which had smaller, and very similar waveforms (Figure 5B). Here waveform based clustering alone yields a very poor result, with a substantial improvement when spike locations are included. This situation is quite typical in recordings from the retina, and, as will be shown below, can lead to wrong assignments even when more advanced methods for waveform-based spike sorting are used.

**Figure 5:**
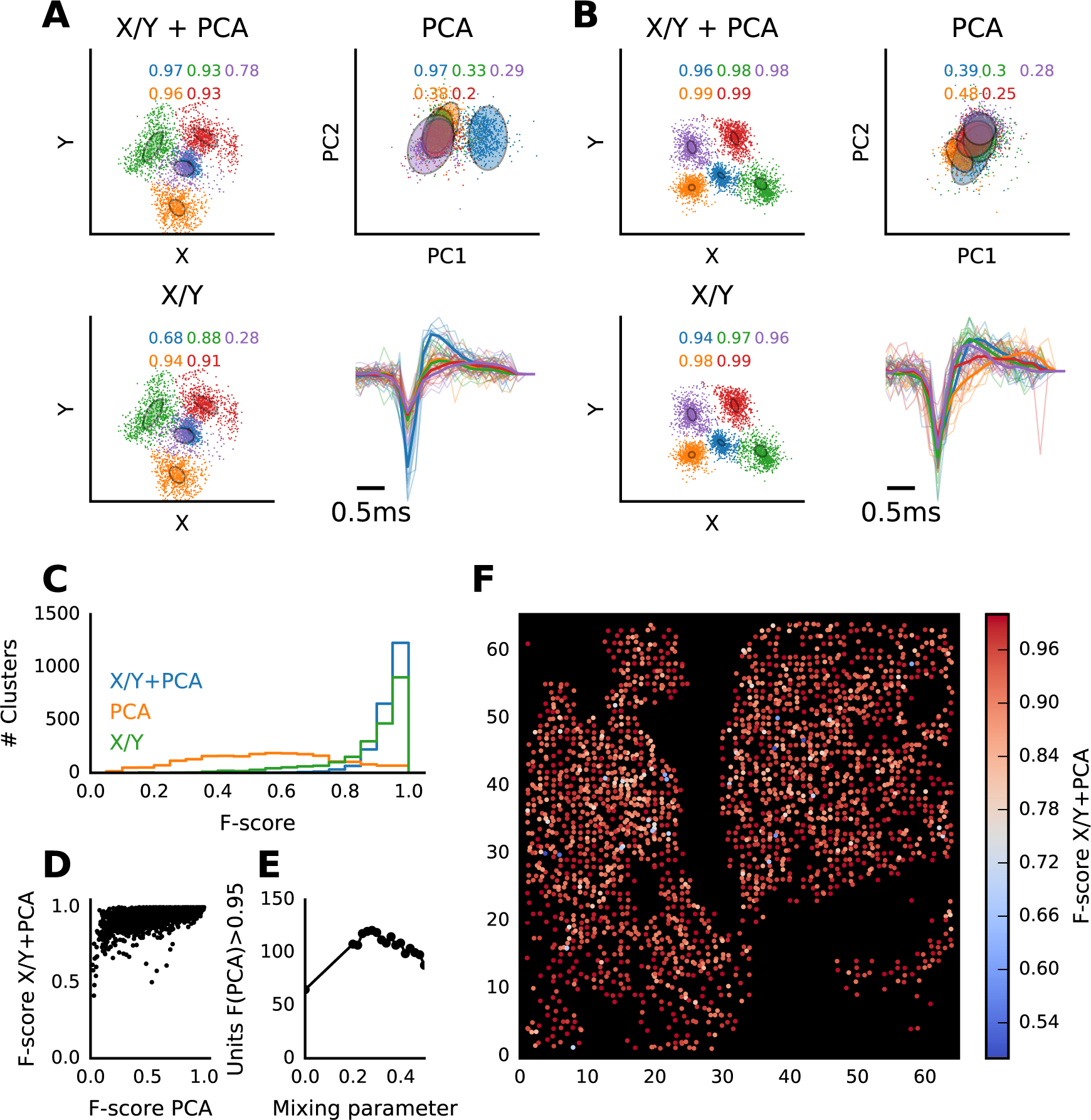
Quantitative assessment of sorting quality with Gaussian mixture models (GMM). (A,B) Two examples of GMMs fit to groups of neighboring units. In both cases, all units within a radius of 42 *μ*m around the unit colored blue were included in the model. The model was then fit to combined spike locations and waveforms (X/Y+PCA), waveforms alone (PCA) or locations alone (X/Y). Spikes are colored to indicate the original cluster assignments. Numbers in each panel are the F-scores for each unit, indicating the average number of false positives and negatives between the two assignments. Each panel also shows examples of spike waveforms and the unit average (thick line), using the same color scheme. (C) Histogram of F-scores of all units in one recording, computed as in A,B. (D) Relationship between F-scores evaluated from waveforms alone and the combined features. (E) Number of units with an F-score>0.95, evaluated from waveforms alone, for different values of the shape mixing parameter *α*. The best overlap is obtained for *α*0.28, the value used in the other examples in this paper. (F) Spatial distribution of F-scores for all units.

The results of an analysis performed on a data set with 7.6 million spikes are summarized in Figure 5C-F. Each of the 2,234 units with a spike rate of at least 0.3 Hz took, in turn, the role of the blue unit in the examples above, for which all units within a radius of 42 *μ*m (or at least the nearest unit) were combined into mixture models. When location and waveform features were used for quality control, 55% of the units (1230) had an F-score>0.95, and 15% (334 units) an F-score>0.99 (Figure 5C, X/Y+PCA). These fractions decreased only slightly when locations (X/Y) were used on their own, but they decreased substantially for waveforms (PCA) alone. A comparison of the F-scores for waveforms and the combined features shows that adding locations improve fits in virtually all cases, but poor scores for waveforms are also more likely to result in lower combined scores (Figure 5D). An inspection of the waveform fits for different mixing coefficients shows there is an optimal value of about *α* = 0.28 (for recordings at 7 kHz), where the number of reliably distinguishable units based on waveforms is maximized (Figure 5E). A spatial overview of these results showed that, as expected, units with low F-scores are primarily found in areas with dense activity (Figure 5F).

### Functional assessment of single unit activity

Our recordings reflect RGC responses to a series of full fleld flashes. We therefore evaluated whether individual sorted units also exhibit the typical On, Off or On-Off responses to light of RGCs. Spike locations, spike waveforms, raster plots and peri-stimulus time histograms of all units in a small patch of retina are shown in Figure 6A,B. These examples demonstrate a clean separation into fast and slow On and Off cells, and neurons with On-Off responses. Importantly, nearby or spatially overlapping neurons generally exhibit unique, different responses, as expected from the mosaic functional organization of RGCs, and there was no apparent mixing of functional properties as would be expected for poorly isolated units.

**Figure 6:**
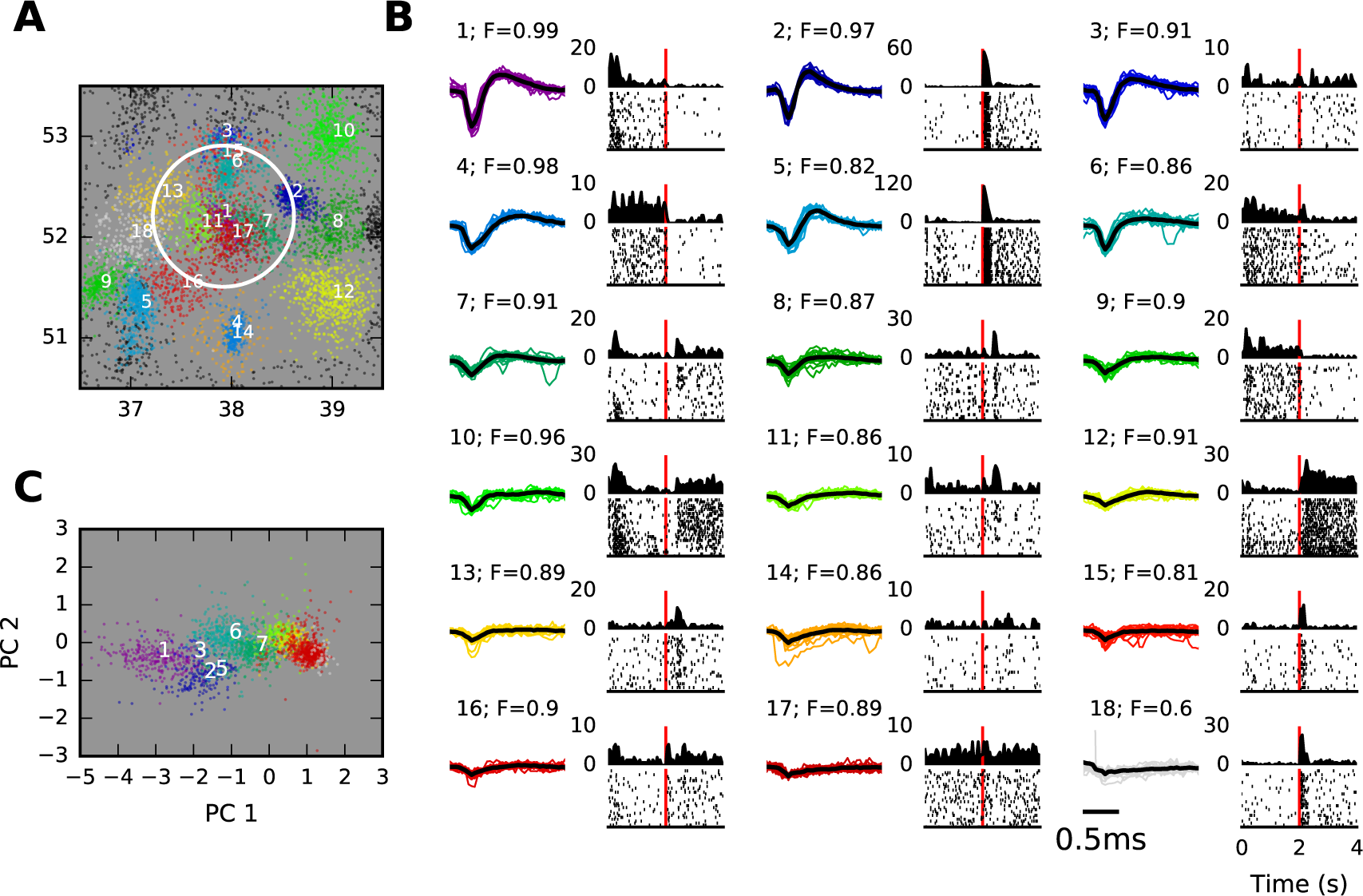
Functional characterization of spike sorted retinal ganglion cells. (A) Spatial locations of individual spikes within a small area on the MEA. Only a subset of spikes are shown for clarity. This area contained 18 units, and unit membership is indicated by color. Spikes of units centered outside the visible area are shown as black dots. Coordinates are in units of electrode distance (42 μm). (B) Overview of the units highlighted in part A, using the same color scheme. Each panel shows example waveforms, the average spike waveform (black line), and the raster and PSTH for full fleld stimulation (2 s bright, 2 dark, red lines indicate stimulus offset time). The unit number and cluster F-score are given above the spike waveforms. (C) Spikes in the circled area in part A, with identical color coding, shown in the space of waveform principal components (PCA space). The same data as in Figure 1 are shown, acquisition rate was 24 kHz.

Notably, this area contained units with vastly different signal strengths. The fact that the majority of these units, also those with very small waveforms, exhibit consistent light responses, shows that much of the signal variance is due to physiological events, and not electrical noise (Muthmann et al., 2015). Yet, as suggested above, units with weak signals may have an incomplete spike record of a neuron, and their clustering may be more ambiguous. Units with well defined waveforms are typically also well separated in their PCA projections, while small waveforms are mainly clustered based on spatial locations (compare units 1-3 with units 5-7 in Figure 6C). As a result, the cluster F-scores (shown above waveforms in Figure 6B) are indeed lower for units with small waveforms, hence a selection of units based on this measure is a reliable way to restrict further analysis to well isolated neurons.

### Comparison with conventional spike sorting

Conventional spike sorting relies entirely on differences in spike waveforms. Our results indicate that this may not be sufficient to obtain good isolation in many cases, yet we only used a two-dimensional representation of the waveforms, which may be too compact to cluster spikes reliably. To evaluate in more detail how the two approaches differ, we compared our method with the outcome of manually curated spike sorting, done on each MEA channel separately. Conventional spike sorting was performed using T-Distribution E-M clustering (Shoham et al., 2003), followed by manual inspection and correction (Plexon Offline Sorter, Dallas, TX).

The data used for this comparison was recorded with 1,024 electrodes at 24 kHz sampling rate (cf. Figure 1), and 538 cluster with at least 200 spikes each were included in the comparison. For each cluster, we located the most similar sorted unit using spike count cross-correlation following binning (each unit is typically found on multiple electrodes), and obtained the number of spikes in the sorted unit that were not part of the cluster (false negatives), and the number of spikes in the cluster not present in the sorted unit (false positives). As for the mixture model above, we then computed precision, recall, and the F-score for each cluster (see Methods).

The three examples in Figure 7A illustrate two common cases we encountered. The first example shows an almost identical assignment for the two methods. This was found for about 18% of the clusters (96 clusters), for which the comparison gave an F-score larger than 0.95 (Figure 7B). Such pairs had very small numbers of false positives and negatives, likely comfortably within the expected variance of the classifier. The pair in Figure 7A, top panel, for instance had 9 false negatives, and no false positives, out of 1818 spikes.

**Figure 7:**
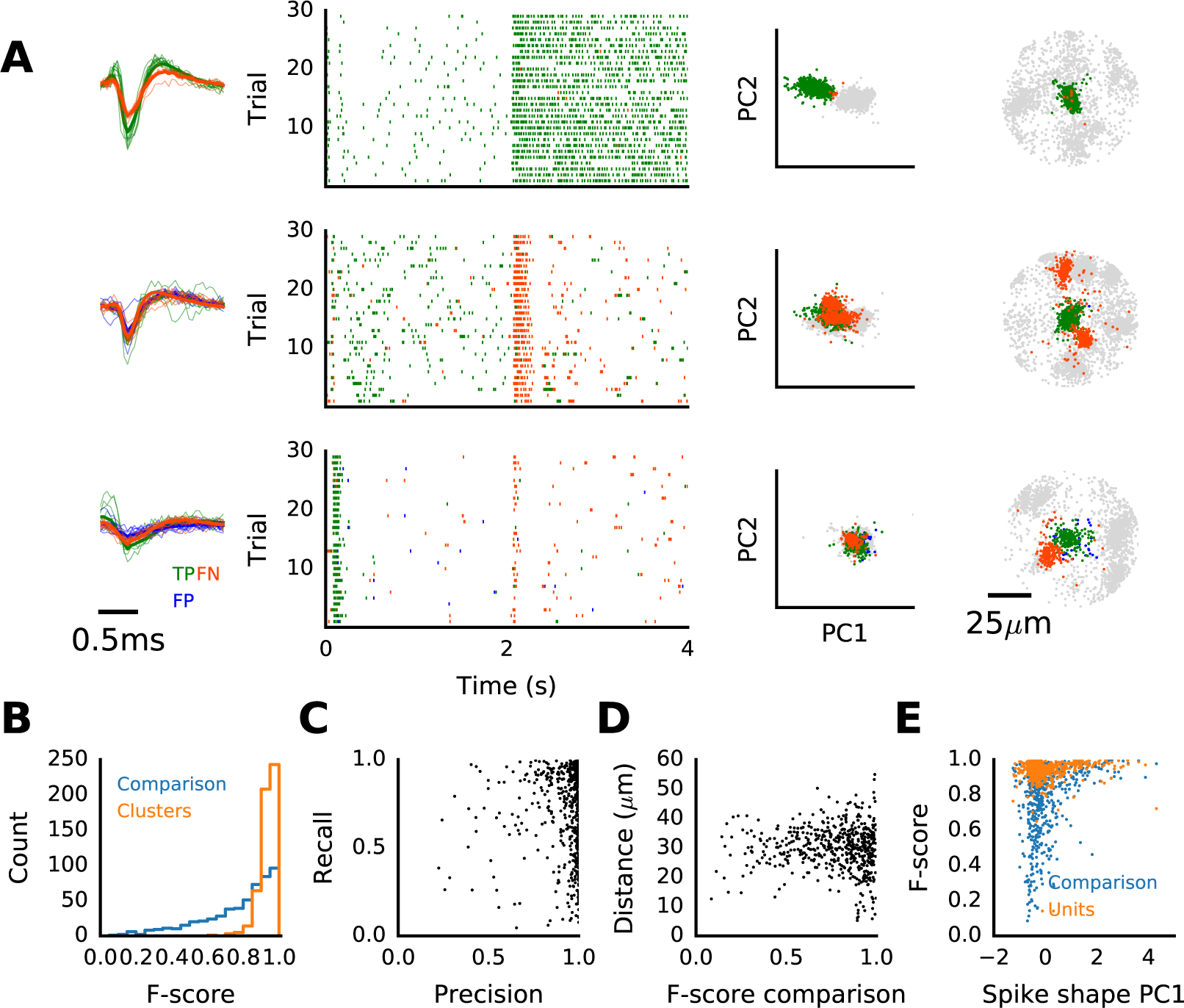
Failure of conventional spike sorting in isolating single units. (A) Examples of three units clustered with our method, compared with corresponding units obtained from con-ventional, spike-shape based sorting. Raster plots show responses to full field flashes (left; 2 s bright, 2 s dark), principal component projections of all spikes found in the area within a radius of 78 *μ*m around the cluster center (middle) and all spikes plotted at their locations (right). Spikes colored green were found in both units, those in orange only in the sorted unit, and those in blue only in the clustered unit. (B) Histograms of F-scores for the comparison (blue) and for mixture model fits for the sorted units (orange). (C) Precision and recall for the comparison, illustrating that low F-scores are primarily due to spikes missing in the clustered unit (orange events in A). (D) Average distance of spikes not included in the clustered unit, measured from the cluster centroid. (E) Comparison of F-scores with the average projection of the waveforms along the first principal component, shown for the comparison of sorting method (blue), and for the mixture model fits of clustered units (orange). The same data as in Figure 1 was used.

For many of the remaining clusters, the F-score was dominated by a sizable fraction of false negatives, spikes in the sorted unit that were not included in the corresponding cluster (units with low recall in Figure 7C). An inspection of the spatial locations of these events showed that false negatives were often located far away from the cluster centroid, and visually appear to be part of another unit (Figure 7A, middle and bottom panels, orange events). The two examples in Figure 7A (middle and bottom panels) show cases where the erroneous assignment by conventional spike sorting has the rather severe consequence of classifying these cells as On-Off rather than On cells.

We found that the inclusion of distant spikes happened frequently, with an average distance of false negatives from the cluster centroid typically around 30 *μ*m (Figure 7D). This suggested they were wrongly included in a sorted unit based on waveform similarities. To see whether these failures are associated with specific waveform features, we compared the F-scores for the comparison with the average projections of the waveforms along their first principal component (PC1, Figure 7E). The PC1 projection provides an indicator of signal quality for each unit (Figure 2B), and indeed lower F-scores were observed almost exclusively for low-scoring units. Hence we conclude that conventional spike sorting only allows reliably isolation of units with strong, very prominent waveform features, while smaller, less distinct waveforms cannot be separated reliably on the exclusive basis of their shape.

## Discussion

Spike sorting is a critical step in the analysis of extracellular electrophysiological recordings. The erroneous assignment of spikes can have rather severe implications for the interpretation of multi neuron activity, so much so that it has been suggested joint models of spike waveforms and neural activity may be required to avoid spurious or biased correlation estimates (Ventura & Gerkin, 2012). In high density recordings, which are increasingly used both for *in vitro* and *in vivo* studies, an additional complication is that the assignment of spikes to single units is a problem with exponential complexity in the number of events, hence requires approximate solutions. Moreover, the sheer size of the data prevents detailed manual inspection and quality control.

Here we solve this task by creating a very efficient data representation, based on spatial spike locations and the most prominent waveform features, which yields a low-dimensional event description and can be clustered efficiently. We developed and tested the algorithm on data acquired with a 4,096 channel MEA with 42 *μ*m electrode pitch, which is sufficient to obtain signals from single neurons in multiple channels. In all examples shown here, we performed clustering in four dimensions, with two dimensions representing waveform features. Adding further features, or using alternative methods for feature extraction, did not yield an increase in performance (not illustrated), which is due to a limited waveform variability in our recordings. This is likely due to the fact that the signals reliably measured with a dense MEA mainly originate from strong currents at the AIS of each neuron, with limited variability between neuron. This, in turn, makes it possible to resolve their spatial origin, but limits the additional amount of information that can be gathered from waveforms. Comparison of optogenetically evoked spikes with anatomical images indeed indicate that detected spikes appear to cluster near the AIS, and that localization alone is sufficiently precise to reliably isolate some units neurons even without using additional waveform features.

Out method can be adjusted to work with other arrays, and potentially also to other probe geometries, as long as a quantity that serves as a location estimate can be obtained reliably. The dimensionality of the clustering step can then be adjusted to exploit higher waveform variability, for instance in *in* vivo recordings. The complexity of the clustering algorithm scales quadratic with the number of spikes, and the highly optimized version used here has a much better performance in situations with prominent spatial clustering. We developed a parallelized implementation, which allows sorting of millions of spikes in minutes (10 million spikes are sorted in about 8 minutes on a 12 core 2.6 GHz Xeon workstation). Together with a method for quality control, this makes it possible to perform parameter sweeps to identify the optimal parameters of the clustering algorithm. Clustering is followed by an automated assessment of clustering quality, based on an ideal (under some assumptions) model of these data. This permits the automated rejection of poorly isolated units, manual inspection of borderline cases. Finally, the data can be inspected using a visualization tool, where further annotation can be performed.

The complete workflow consists of first performing event detection, followed by spatial localization, clustering, quality control, and finally an optional manual inspection. The former two are described in detail in Muthmann et al. (2015), and currently constitute the main bottleneck of the analysis chain. Detection currently performs about 1/4x real time, and scales linearly with recording duration. The complexity of the spike localization scales linearly with the number of detected events, and runs roughly in real time for recordings with normal spike rates. Both methods are parallelizable, and implementations are under development.

Importantly, working with high density recordings during this study has revealed significant limitations of purely shape-based spike sorting. A major issue is that it can be hard or even impossible to decide how many units the signal from a single electrode contains. If an electrode is positioned close to a group of neurons, one or perhaps two units with strong signals may have sufficiently distinct waveforms to be separable. Yet comparison with spike locations showed that weaker signals with different spatial origin are generally not distinguishable based on shape alone. As illustrated, we frequently found cases where spikes of neurons with entirely different physiological signatures were mixed by shape-based sorting, a problem that cannot be avoided even by careful manual inspection. In contrast, our method can cope well with such situations because spatial location estimates are sufficiently precise to disambiguate such cases. A main factor affecting sorting performance is thus the noise and bias in spatial localization, which both depend on the signal quality (Muthmann et al., 2015).

To achieve good sorting quality, we found that a four-dimensional representation of spikes, which includes the two main principal components of the averaged waveforms, was sufficient to obtain highly satisfactory results, even at low acquisition rates of 7 kHz. Adding more dimensions could yield further improvements, but also affected the performance of the clustering algorithm. Future work could thus focus on an improved representation of the most important spike waveform features.

A different strategy for high density recordings, outlined by Marre et al. (2012), is to estimate spatio-temporal templates, which are then used to identify spikes from each neuron (see also Dragas et al., 2014). This shifts the computational burden from spatial interpolation and source localization in our method to the deconvolution of spikes from raw data. We found that adding additional shape criteria in the detection stage could lead to false negatives, suggesting that templates will only yield reliable results if the firing rate of the neurons is high enough so they can be estimated well enough. A third method, recently developed by Rossant et al. (2016) for high density *in vivo* probes, achieves to reduce complexity by masking out irrelevant parts of the data based on geometric constraints, before fitting a mixture model and clustering the data. This avoids an early discarding of potentially useful information, which our method does by using signal interpolation, and Marre et al. (2012) by creating templates. On the other hand, this method is, while potentially more precise, computationally more demanding, and hence only preferable for data from hundreds of channels.

Large-scale, high-density electrophysiological recording approaches are relatively new, but they are rapidly becoming more widespread, offering unique opportunities to investigate small scale neuronal networks at an exquisite level of details, unraveling important networks properties that could not be detected with fewer and more distant recording probes. However, these exciting developments in network neuroscience come at the expense of new challenges in terms of reliable isolation of signals from single cells that are densely packed with each other. The method we have presented in this study will greatly help alleviating these problems, providing reliable and fast separation of signals originating from thousands of neurons communicating with each other.

## Experimental Procedures

### Electrophysiology

Experimental procedures were approved and carried out in accordance to the guidance provided by the UK Home Office, Animals (Scientific procedures) Act 1986 for experiments on retinas performed at Newcastle University, UK, and by the institutional IIT Ethic Committee and by the Italian Ministry of Health and Animal Care (Authorization ID 227, Prot. 4127 March 25, 2008) for experiments on neural cultures performed at IIT, Genova. Previously acquired data from hippocampal cultures were used, for a detailed description see Panas et al. (2015).

Experiments on the retina were performed on adult wild type mice (C57Bl/6, aged postnatal day (P) 27-39) or on B6.Cg-Tg(Thy1-COP4/EYFP)9Gfng/J mice (Thy1-ChR2-YFP; Jackson Laboratories, Bar Harbor, USA; RRID:IMSR_JAX:007615) aged P69-96. In these mice, thy1-expressing RGCs coexpress ChR2 and yellow fluorescent protein, allowing visualization under fluorescence microscopy. High density recordings from the RGC layer were performed using the BioCam4096 platform with APS MEA chips type BioChip 4096S (3Brain GmbH, Switzerland), providing 4096 square microelectrodes (21 μm x 21 μm) on an active area of 2.67 mm x 2.67 mm, aligned in a square grid with 42 μm spacing. The platform records at a sampling rate of about 7 kHz/electrode when measuring from the full 64 x 64 electrode array, and was also reconfigured to sample at 24 kHz when recording from one quarter of all electrodes. Raw data were visualized and recorded with the BrainWave software provided with the BioCam4096 platform. Activity was recorded at 12 bits resolution per electrode, low-pass filtered at 5 kHz with the on-chip filter and high-pass filtered by setting the digital high-pass filter of the platform at 0.1 Hz.

Mice were killed by cervical dislocation and enucleated prior to retinal isolation. The isolated retina was placed, RGC layer facing down, onto the MEA. Coupling between the tissue and the electrodes was achieved by fiattening the retina on the array under a small piece of polyester membrane filter (Sterlitech, Kent, WA) maintained in place by a stainless steel ring. The retina was kept at 32 °C with an in-line heather (Warner Instruments) and continuously perfused using a peristaltic pump (~1 ml/min) with artificial cerebrospinal fluid (aCSF) containing the following (in mM): 118 NaCl, 25 NaHCO_3_, 1 NaH_2_ PO_4_, 3 KCl, 1 MgCl_2_, 2 CaCl_2_, and 10 glucose, equilibrated with 95 % O_2_ and 5 % CO_2_. All preparations were performed under dim red light and the room was maintained in darkness throughout the experiment.

### Visual and optogenetic stimulation

Visual stimuli (664x664 pixel images for a total area of 2.67x2.67 mm) were presented using a custom built high-resolution photostimulation system based on a DLP video projector (lightCrafter, Texas Instruments, USA) combined with a custom made photostimulation software and synchronized with the recording system (Portelli et al., 2016). Neutral density filters (4.5 - 1.9) were used to control the amount of light falling on the retina.

Photoreceptor-driven responses were acquired at a maximum irradiance of 4 *μ*W/cm^2^ (ND 4.5), low enough to avoid eliciting ChR2-driven responses in the ChR2 retinas. To isolate ChR2 responses from photoreceptor-driven responses in these same retinas, we decreased synaptic transmission by increasing the MgCl_2_ concentration to 2.5mM and by decreasing the CaCl_2_ concentration to 0.5mM in the aCSF solution, and used 20 *μ*m DNQX, and 20 *μ*m L-AP4 (Tocris Bioscience, UK) to respectively block metabotropic ionotropic and activate metabotropic (MGLUR6) glutamate receptors, to block all responses to light in bipolar cells and their postsynaptic RGC partners. We used the broad RGB spectrum of the DLP projector with a maximum irradiance of 0.87 mW/cm^2^ (ND 2.2) to evoke ChR2 responses. A battery of conventional stimuli (full field flashes, moving gratings) and spatio-temporal white noise were used, and responses to used repetitive (30x) full field stimuli (0.5 Hz) were analyzed in Figures 6 and 7.

### Spike detection, localization and selection

The procedures for spike detection and current source localization were described in detail elsewhere (Muthmann et al., 2015). First, weighted interpolated signals were generated using two spatial templates to capture both spikes originating either close to or between electrodes. A five channel template with a strong relative weight for the central electrode and weaker weights for the four surrounding electrodes emphasized current sources close to electrodes. Sources between electrodes were captured by a four channel template, which generated a signal that was added to the data as a virtual channel. A running estimate of the signal baseline and noise level was computed from percentiles for each signal, and putative spikes were detected as threshold crossings. This procedure ensures that temporally overlapping spikes are detected as long as they leave a distinct spatial footprint. Next, source locations were estimated for each event by considering the spatial signal spread over neighboring electrodes. Briefly, all signals were baseline-subtracted and inverted, then the median signal was subtracted to minimize bias due to noise, the signal clipped to positive values, and the center of mass determined.

To filter out noise and poorly detected neurons in recordings at low sampling rates (7 kHz), we implemented an automated post hoc rejection of events for recordings at lower sampling rates. To this end, noise events were sampled from areas on the MEA where no activity was recorded, such as at incisions or uncovered areas (identifiable by low spike counts). Up to 1000 of such events, as well as up to 1000 events with large amplitudes were used to train a Support Vector Machine with radial basis functions. This model was then used to classify events as true spikes or noise (see Supplemental Information, Figure S1).

### Spike clustering

Data points were clustered together using an implementation of the Mean Shift algorithm (Comaniciu & Meer, 2002) available in the scikit-learn open source machine learning library (Pedregosa et al., 2011). Importantly, this algorithm did not require the knowledge of the desired number of clusters; it depended, instead, on a single parameter, the bandwidth *h*, which determined the expected cluster size. This size can be estimated from a typical spatial cluster size in an activity plot (Figure 1B), and was here set to 12.6 *μ*m (the average width of clusters). To combine spatial and waveform information, the clustering process was run on a four-dimensional space consisting of two dimensions indicating the location of each event on the chip, *x* and *y*, and two dimensions representing the first two principal components of the event’s waveform. The latter were multiplied by an additional dimensional constant *α* that tuned the relative importance of the waveform components compared to the spatial coordinates. The optimal value of *α* varies slightly for different data sets, and is typically in the range 0.28-0.34 (Figure 5). To parallelize this algorithm, we exploited the fact that all points follow a local density gradient until they converge to a local maximum, the center of a cluster. Because every data point does so independently of the others, this process is run in parallel, which improved performance roughly proportionally to the number of available CPUs. The relevant code has been merged into the scikit-learn Python library.

### Quality metric

Following Hill et al. (2011), we fitted a multivariate Gaussian mixture model to a set of *N* clusters, then estimated cluster overlap using posterior probabilities to obtain the probability of incorrect assignments under the assumption of a Gaussian cluster shape. The model is fit in six dimensions, with the two spatial coordinates and the projections of the spike waveform along the first four principal components. For each cluster, we assume that only spikes in nearby clusters interfere with the sorting. Therefore, all clusters within a radius of 42 *μ*m (electrode pitch) are included in the model, or at least the closest neighbor if no cluster was found within this area. To obtain meaningful fits for sets of clusters with a very unbalanced number of spikes, first a Gaussian is fit to each cluster individually, which are then combined into a mixture model.

The quality of the assignment is then evaluated as follows. Let the probability that spike *s* is in cluster *c* be *P* (*C* = *c*|*S* = *s*): the,estimated fraction of spikes in cluster *k* that could belong to cluster *i* is given by 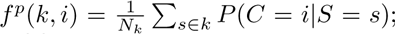 by generalizing to all other clusters we obtained the number of false positives in *k*:

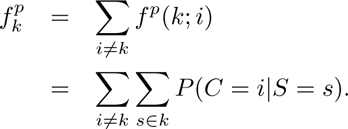

Correspondingly, we could estimate the fraction of spikes in cluster *c* that was expected to be assigned to other (i.e. wrong) clusters and obtained the number of false negatives as:

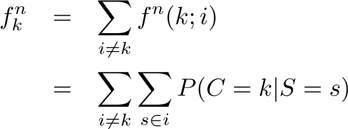

The probabilities *P*(*C* = *c, S* = *s*) were given by mixture model. To obtain a single quality measure, we compute precision (*P*_*k*_) and recall (*R*_*k*_):

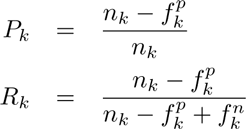

The harmonic mean of these quantifies yields the F-score:

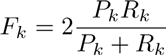

### Confocal imaging and image analysis

To achieve a precise alignment of RGCs with recording electrodes, the retina had to be imaged on chip with photoreceptors facing upwards. As reported before, we found that photoreceptors appear to act as microlenses guiding and scattering the incident excitation light (Denk & Detwiler, 1999). We circumvented this problem by embedding the fixed tissue in a mounting medium with matching refractive index. After the recording, the retina was immediately fixed with 4% paraformaldehyde (in 0.1 M PBS, 200 mM sucrose) on the MEA chip for 1 hour. In preliminary experiments we determined that tissue shrinkage, which may interfere with activity alignment, is negligible for this protocol. The retina was then rinsed several times with 0.1 M PBS and embedded with Vectashield (Vector Laboratories, UK) and sealed with a coverslip (Menzel Glaeser, Germany). Imaging was performed with a Leica SP5 confocal upright microscope supplied with a 25x / 0.95NA WD 2.5 mm water immersion objective for an optimal signal collection focusing on 8x8 electrode arrays in 300x300 *μ*m field of view. In each field, images (2048x2048 pixels) were acquired in z-stacks in tissue thickness 60-100 *μ*m with optical slicing that corresponded to 30-50 image planes in each tissue volume. Acquisition parameter optimization revealed that a lateral resolution of 200 nm per pixel, just above the diffraction limit, and optical slicing of 550 nm provided an adequate trade-off between the level of image detail for morphological analysis and the acquisition time minimizing the risk of photo damage for long exposure. Microscope parameter optimization was performed using tools to increase the signal-to-noise ratio, including high number of frame averaging with an upper limit determined by safe levels of laser power to protect the tissue, and post image processing methods using deconvolution. The degree of image blurring of the microscope can be characterized by the point spread function (PSF) in image formation theory. When the explicit knowledge of the experimental PSF is unknown, which is quite challenging to determine in thick volume of tissues with high inhomogeneities in different depth locations for 3D reconstruction, blind deconvolution algorithms are extensively used in image restoration. The Richardson-Lucy (RL) (Richardson, 1972; Lucy, 1974) was used, with the number of iterations optimized for maximum image sharpness, maximum contrast and minimum image distortion, depending on the noise level and image blur, usually in the range from 3 to 10. In addition to the fluorescence signals in specific fields, large-field images including images of the electrode array were also acquired in order to enable the co-localization of images with RGC spiking activity.

In one Thy1 YFP/ChR2 retina, RGC somata were manually annotated in selected subfields where activity was recorded, and the confocal images of the RGC layer were spatially aligned with the estimated locations of detected events. To this end, the active area of one electrode was determined, and the remaining electrode locations were computed generating a regular grid using the 42 *μ*m electrode spacing. The images and soma locations were then transformed into array coordinates, and spike locations were overlaid with the retinal image.

## Author Contributions

Conceptualization, G.H., M.S., E.S., and M.H.H.; Methodology, G.H, M.S., J-O.M., S.P., I.E.K., S.U., E.S., and M.H.H.; Software, M.S., S.P., J-O.M., C.J.R., A.P.E. and M.H.H.; Formal Analysis, M.S., S.U., and M.H.H.; Investigation, G.H., M.S., S.P., I.E.K., and S.U.; Resources, A.M., and L.B.; Data Curation, G.H., A.M., L.B., and E.S.; Writing - Original Draft, M.H.H. with input from co-authors; Supervision, L.B., V.M., D.S., F.C.Z, E.S., and M.H.H.; Funding Acquisition, L.B., V.M., D.S., F.C.Z., E.S., and M.H.H.

## Acknowledgements

We thank Fernando Rozenblit, Vidhyasankar Krishnamoorthy and Amos Storkey for valuable input. This work was supported by the 7th Framework Program for Research of the European Commission (Grant agreement 600847: RENVISION, project of the Future and Emerging Technologies (FET) program Neuro-bio-inspired systems FET-Proactive Initiative) and the Wellcome Trust (grant number 096975/Z/11/Z). M.S. was supported by the EuroSPIN Erasmus Mundus programme and the EPSRC Doctoral Training Centre in Neuroinformatics (EP/F500385/1 and BB/F529254/1), and J-O.M. by the EuroSPIN Erasmus Mundus programme and by NCBS/TIFR.

